# Species-habitat networks: Bridging applied ecology and network theory

**DOI:** 10.1101/326041

**Authors:** Lorenzo Marini, Ignasi Bartomeus, Romina Rader, Francesco Lami

**Affiliations:** DAFNAE, University of Padova, Viale dell’Università 16, 35020 Legnaro, Padova, Italy; Dpto. Ecologia Integrativa, Estacion Biologica de Dõnana (EBD-CSIC), Avda. Americo Vespucio 26, Isla de la Cartuja, 41092 Sevilla, Spain; Ecosystem Management, School of Environment and Rural Sciences, University of New England, Armidale, NSW 2351, Australia

**Keywords:** Bipartite networks, Habitat use, Landscape heterogeneity, Land-use change, Patch-mosaic, Stability

## Abstract

Land-use change is massively reshaping terrestrial ecosystems worldwide, and is recognized as a key driver of biodiversity loss with negative consequences on ecosystem functioning. Understanding how species use resources across landscapes is essential for the design of effective management strategies. Despite recent advances in theoretical ecology, there is still a gap between theory and applied ecological science and we lack the tools to manage entire landscapes to maximize biodiversity conservation and ecosystem service delivery. Here, we propose a new approach that uses existing bipartite networks to create species-habitat networks. Networks enable powerful visualizations via a common language that defines most processes in terms of nodes and links. This approach explicitly links multiple species and habitat resources, provides tools to estimate the importance of particular species in a given landscape, and quantifies emerging properties of entire habitat networks. Most existing metrics used to study properties of bipartite ecological networks can easily be adapted to investigate species-habitat relationships. One key advantage of this approach is that the scale of the derived ecological information will match the scale of management interventions. The flexibility of the proposed approach is that it can be easily applied across a range of ecological fields such as species conservation, habitat restoration, ecosystem services management, or invasion ecology. Network emerging properties could also be used to test the effects of large scale drivers of global change upon ecosystem structure and stability.

## Community ecology across heterogeneous landscapes

Understanding how species use resources across landscapes is essential for the design of effective management strategies to support biodiversity and ecosystems services. By using conceptual or mathematical models, theoretical ecology has greatly improved our understanding of the dynamic principles which govern the way populations and communities respond to landscape processes (Forman 1995, MacArthur and Wilson 2001, Loreau et al. 2003). To date, patch-matrix models rooted in meta-community (Leibold et al. 2004) or island biogeography theory have largely focused on species responses to the amount and configuration of remnant habitats within a hostile matrix (Tscharntke et al. 2012, Hadley and Betts 2016) (Fig. 1A). Central tenets of these models are that species dispersal (i.e. the flow of individuals) occurs mainly between patches and that the focal population mostly relies on resources occurring within a specific habitat. As it is becoming increasingly clear that many species utilize a range of different habitats of varying qualities (Ricketts 2001, Tews et al. 2004, Driscoll et al. 2013), landscape ecology has moved beyond the dichotomy of patch-matrix models to explicitly incorporate landscape heterogeneity (Wiens et al. 1993, Fischer and Lindenmayer 2006, Cushman et al. 2010, Brudvig et al. 2017). Yet, while the meta-ecosystem concept has provided fundamental insights into the dynamics and functioning of ecosystems from local to regional scales (Loreau et al. 2003), there is still a gap between theory and empirical research and few methods have linked species and habitats in real landscapes (Gounand et al. 2017)

**Figure 1.**
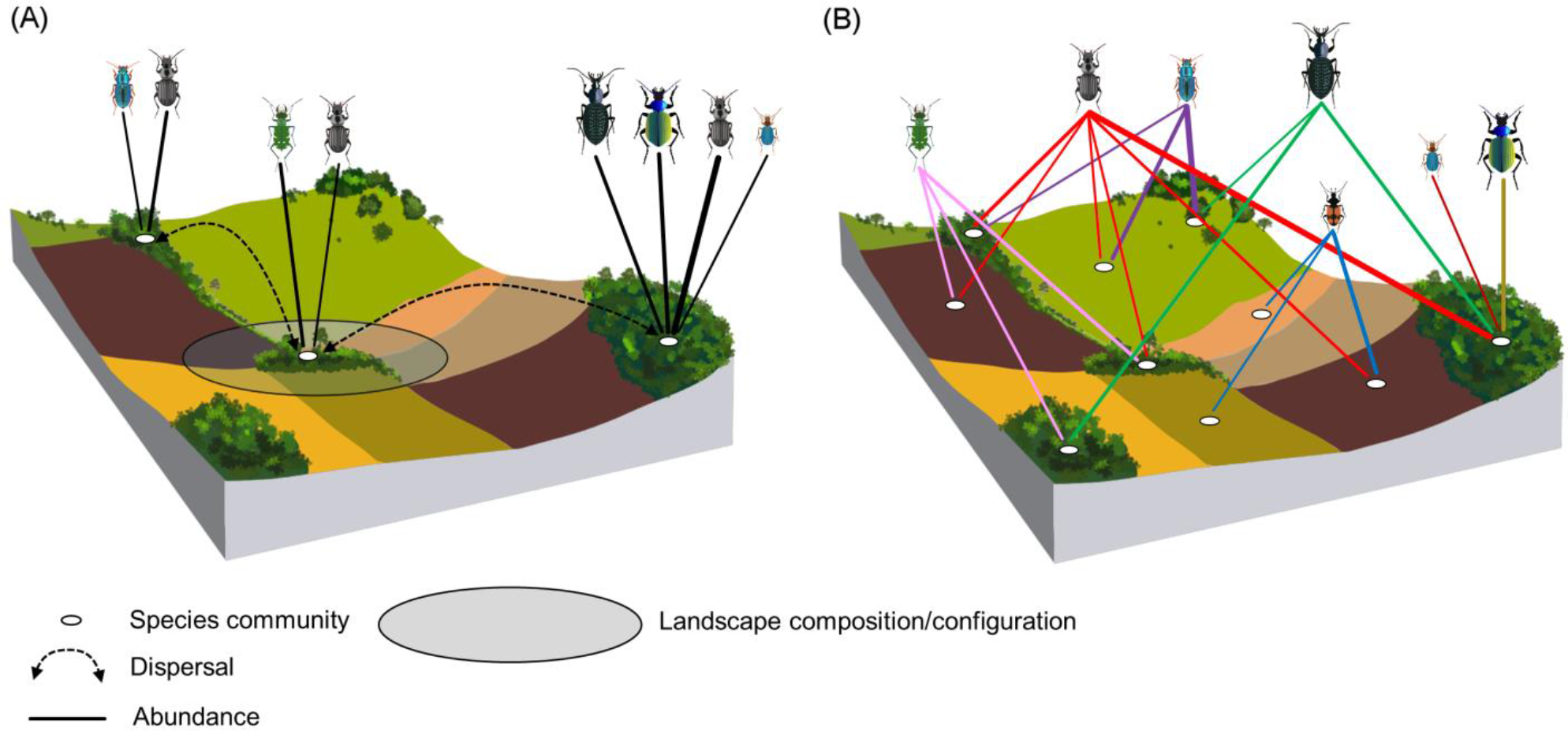
(A) Current spatial approach to study species community dynamics across heterogeneous landscapes. Meta-community ecology focuses on patches of one focal habitat embedded in a homogenous hostile matrix and linked through dispersal (black dotted arrows). Most empirical research in landscape ecology focuses on local habitat vs. landscape mosaic, where the landscape is quantified in terms of composition and/or configuration around a central point where the community is sampled (shadow buffer). Neither approach accounts for the interactions between multiple species and habitats outside the focal habitat. (B) The species-habitat network whereby the whole landscape is sampled and the species are quantified at multiple sites (line width proportional to species abundance). The landscape can be classified in patches according to the functional role of the different habitats for the target species community. If species do not occupy readily identifiable habitat patches, a continuous variation in habitat quality and available resources around the sampling sites can replace the discrete habitat categorization.

The field of landscape ecology has made significant inroads toward understanding community responses to landscape processes at multiple spatial scales (Turner 2005, Fahrig et al. 2011). This empirical research has driven the field of applied ecology forward by providing a solid evidence base for managers and policy makers (Tscharntke et al. 2005, 2012, Mayer et al. 2016). However, most of these studies are based on another dichotomy, i.e. a focal local habitat vs. the surrounding landscape. Often the species community of interest is only sampled in one habitat and related to the landscape by using the proportion of suitable or unsuitable habitats (Fig. 1A). When landscape heterogeneity is taken into account, it is usually quantified using metrics that collapse complex processes into single indices (Frazier and Kedron 2017). Many examples of this approach in applied ecology exist (Clough et al. 2014) and recent advances in ecosystem services research have successfully applied the same approach to study key functions such as seed dispersal, biocontrol (Schellhorn et al. 2015) or pollination (Kennedy et al. 2013). One downside of this research, however, is the lack of a mechanistic understanding of the links between multiple habitats and community-level processes, indicating the need for broader conceptual frameworks of spatial patterns.

Network approaches based on graph theory have been increasingly applied to the problem of describing complex and dynamic community level changes in ecology (Bascompte and Jordano 2007, Minor and Urban 2008, Memmott 2009, Blonder et al. 2012, Burkle et al. 2013, Albert et al. 2017, Gilarranz et al. 2017, Harvey et al. 2017). The network paradigm is based on the representation of emerging properties of studied systems as oriented graphs: any system is traced back to a set of nodes (its constituent units) linked by edges corresponding to the relationships between nodes. This allows for a straightforward quantitative formalization of systems by computing mathematical descriptors of such graphs. In this way, network tools have already been applied to elucidate landscape processes, i.e. habitat patches have been represented as nodes and linked via dispersal to model connectivity at multiple spatial scales (Burns and Zotz 2010, Dale and Fortin 2010, Gonzalez et al. 2011). While these pioneering approaches have enabled the link between habitat configuration and species dispersal, they have failed both to upscale from species to community level, and to consider multiple habitat types simultaneously.

## Beyond the focal habitat: Introducing the species-habitat network

Traditionally, patch-mosaic models have defined landscapes as complex and heterogeneous mosaics, constituted of many interacting discrete habitat patches. More recently, several gradient models of landscape structure have challenged the mosaic paradigm (Fischer and Lindenmayer 2006), suggesting that landscape heterogeneity should be modelled using multiple, continuous environmental gradients (Cushman et al. 2010). In both cases, explicitly accounting for species resource use requires the sampling of target species in multiple sites across the landscape. These ideas have led us to consider the whole landscape as a unit to quantify and analyse community response to landscape processes (Fig. 1B). Integrating and analysing species use of multiple sites within a landscape may seem a daunting task, especially because the number of species-sites links scale exponentially with the number of species and sites sampled. Fortunately, tools developed from ecological network theory can be used to analyse and describe such complex interactions. In particular, we advocate the modelling of species-habitat interactions as bipartite networks (Box 1), analogous to those describing antagonistic or mutualistic interactions (Bascompte and Jordano 2007). Bipartite networks are networks in which two types of nodes exist, and interactions are analysed only between nodes of different types. In the most simple case, habitat types and the species occurring within each habitat constitute the two types of nodes. The links between species and habitats are represented by the number of individuals occurring in a certain habitat at any given moment. The flexibility of the proposed approach allows habitat nodes to be further defined as individual sites where the community was sampled (Burns and Zotz 2010). This definition of a node can incorporate the underlying spatial processes associated with differences in landscape composition and configuration. That is, each individual site could affect network topology and stability depending on its attributes such as habitat quality, disturbance or connectivity. When species do not occupy readily identifiable habitat patches, a continuous variation in habitat quality and available resources around the sampling points can replace a discrete habitat categorization. Once the nodes are defined, the links need to be carefully formulated as they can affect the ecological interpretation of the species-habitat network. The operational definition of a link is the occurrence/abundance of a particular species in a certain location (Box 1). The focal species community would usually belong to the same trophic level sharing a similar functional role. Examples could include lichens, pollinators, ground-dwelling predatory arthropods, insectivorous mammals, etc.

#### Box 1 Sampling a species-habitat network

Any heterogeneous landscape and the species using its resources can be visualized as a bipartite network. In this example, we will consider the butterfly species occurring across an agricultural landscape in a temperate region. In the example, we sample the butterfly species occurring at 15 sites belonging to five habitats (forest, grassland, shrubland, fallow and urban area) within a landscape mosaic (1.5 × 1.5 km) (Figure IA).

**Figure I.**
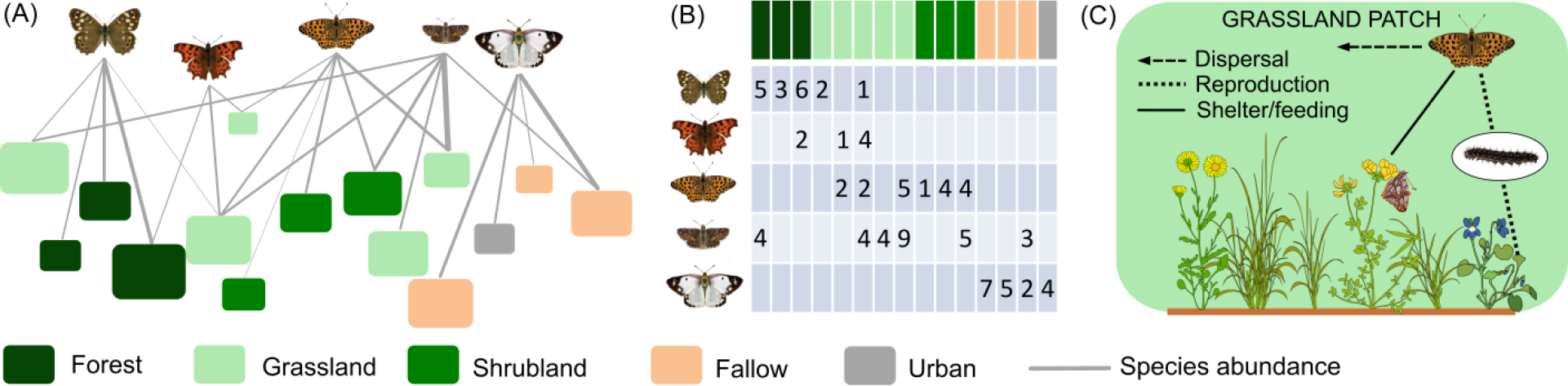
(A) A species-habitat network of 15 sites of varying size and quality belonging to five habitats in which a butterfly community is sampled, (B) data matrix that can be derived from the sampling, and (C) example of types of species-habitat link.

The 15 sites may represent different qualities (e.g. size, tree cover or management) and spatial configurations. If species do not occupy readily identifiable habitat patches (e.g. forest-shrub gradient), a continuous variation in habitat quality and available resources around the sampling points can replace the discrete habitat categorization. The nodes in the network are represented by the butterfly species and the sampling sites. The numbers indicate the link strength (number of individuals in each site) (Figure IB). The number of sampling sites is selected to be representative of the resources and habitat diversity. In the example, the butterfly-habitat network is built using the cumulative abundance from three rounds of sampling (spring, early summer and summer) using a transect walk method. In a transect walk, butterflies are recorded in a fixed width band (typically 5 m wide) within each site. Particular attention should be paid to the functional interpretation of the links. If we consider one grassland in this network (Figure IC), a butterfly species is recorded in that site because individuals can use multiple resources (e.g. host plants for reproduction, nectar for adult feeding or plants for roosting or shelter) or simply because individuals are using that site as a stepping stone for dispersal. Hence, the choice of the sampling method will dictate the interpretation of the ecological data. In this case a transect walk emphasizes the weight of adult feeding over reproduction. On the other hand, an alternative sampling focused on butterfly larvae and host plants can inform about species habitat use for reproduction (Dainese et al. 2017). This idea can be expanded to any taxa that use resources across heterogeneous landscapes.

## Building species-habitat networks

Several studies have shown that individual species and community responses to landscape processes depend on the spatial scale over which the landscape metrics are quantified (Steffan-Dewenter et al. 2002, Fahrig et al. 2011). The selection of the appropriate spatial extent is thus the first key issue that needs to be addressed when building a species-habitat network. The spatial extent in which the community is sampled should be selected according to species’ foraging ranges (e.g. for mobile organisms) or propagule dispersal (e.g. for sessile organisms) and to the ecological hypotheses underpinning the study. This issue is similar to the selection of buffer radii when adopting a traditional approach to quantify landscape composition or configuration. Once the spatial extent is defined, the species communities need to be sampled across the landscape. In most cases, the most pragmatic solution would be to adopt a ‘habitat-centric’ approach where the number of sampled sites is proportional to the habitat area. It is important to stress that as the spatial extent of the habitat mosaic used by the species is generally large (e.g. 1-10 km for mobile organisms), it is likely that most surveyed species-habitat networks would be subunits of much larger networks (Jordano 2016).

While the definition of species as nodes is usually straightforward, the way in which habitats are defined as nodes can be more complex (Frazier and Kedron 2017). Spatial grain and habitat classification can affect the topology (and hence interpretation) of the network. In modified landscapes, different habitats are often organized in patches, which can be defined as discrete areas with a definite shape, size and configuration. The focal species community may be used to guide the identification of habitat types that are functionally relevant. From an operational point of view, we suggest that habitat nodes are defined according to the dominant vegetation (e.g. crop, forest, semi-natural grasslands, etc.), accounting for differences in structure and function for different communities. However, a species-habitat network does not necessarily require a patchy habitat structure and a representation of landscape heterogeneity using continuous gradients can also be incorporated in this framework (Fischer and Lindenmayer 2006).

Finally, understanding how and why the topology of the networks changes over time, and how these changes affect species resource use across the landscape, can help to predict the consequences of human impacts upon community dynamics (Blonder et al. 2012). Incorporating a temporal perspective, however, requires careful thought of the timing (when) and spacing (how frequently) of the sampling. For instance, a longitudinal design with repeated observations within or across years can inform the degree of temporal variability in the species-habitat use (Laliberté and Tylianakis 2010). In the case of species-habitat networks at equilibrium, system stability to perturbations can be further investigated using both empirical and simulation models (May 1972, Memmott et al. 2007, Thébault and Fontaine 2010).

## Use and limitations of the framework

There are several important conditions to note when operationalising species-habitat networks. First, users must ensure that the data inputs are realistic and relevant to the community sampled to ensure meaningful results are obtained through the network analysis. For sessile organisms such as lichen or plant species, occurrence directly links to resource use and habitat preference (Burns and Zotz 2010). On the contrary for mobile organisms that use multiple resources, species occurrence can assume different ecological meanings (Kremen et al. 2007). If we consider a specific habitat, a species can be recorded at that site because individuals can use multiple resources (e.g. host plants for reproduction, preys, nesting site or structure for roosting or shelter) or simply because individuals are using that site as a stepping stone for dispersal. Hence, depending on the species traits and the sampling method chosen the species-habitat networks can capture different community properties (Box 1).

Second, not all taxa can be appropriately described by species-habitat networks. One situation where the framework is unlikely to be applicable is when average species dispersal in the community is too large (e.g. large mammals or birds) compared with the feasibility of field sampling.

Third, the species-habitat networks may be limited in use when the landscape structure is characterized by high habitat heterogeneity at a spatial scale much smaller than the average species dispersal.

For instance, sampling insect communities in highly complex forest landscapes such as those in tropical regions might be challenging. On the contrary, human-altered landscapes with high contrast between habitat types provide ideal conditions to apply the framework.

Fourth, the required sampling effort is likely to be relatively higher than traditional observational landscape studies. However, sampling a greater number of sites will more likely capture the intrinsically high complexity of community response to landscape processes, which is pivotal to adequately address particular ecological questions. Additionally, while species-level is the obvious unit to consider in this context, species may also be grouped using functional traits to reduce network dimensionality (Eklöf et al. 2013). As for most empirical ecological interaction networks, species-habitat networks would suffer to some extent from under-sampling. Hence, limitations imposed by sampling incompleteness need to be carefully explored (Vizentin-Bugoni et al. 2016). Robust estimates of the actual number of individuals of mobile species occurring across a landscape mosaic require an adequate sampling effort that needs to be explicitly evaluated (Jordano 2016).

## Tools for analysing species-habitat networks

The appeal of a network approach is that they enable very powerful visualizations via a common language that defines most processes in terms of nodes and links. Most existing metrics used to study properties of bipartite ecological networks can easily be adapted to the study of species-habitat networks. These metrics can be broadly divided in two groups: emergent properties of the whole network and node-level metrics that measure the role of single nodes (i.e. single habitat sites or species) in the network (Dormann et al. 2009) (Fig. 2). As metric choice will depend on the nature of the question, we advocate a hypothesis-driven approach whereby users decide *a priori* which metrics will address which research question.

**Figure 2.**
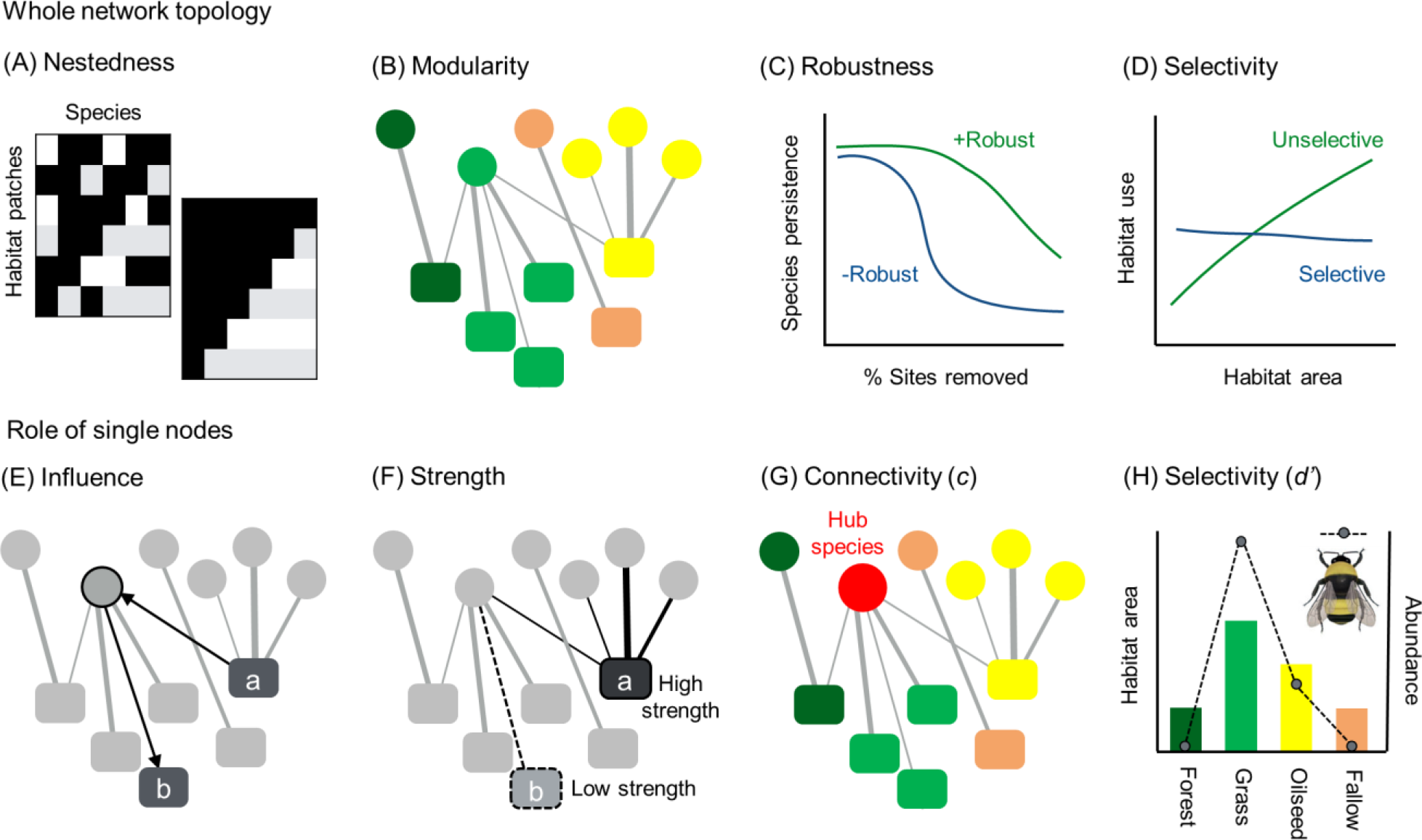
An untapped network toolbox for assessing species-habitat links. Bipartite network analysis is a mature field able to identify emerging properties of a system (A-D) as well as the roles that individual nodes (species or habitat sites) play in the network (E-H). Circles and rectangles represent species and habitat sites, respectively. Here, we present only a few examples of the metrics that can be computed (Blüthgen *et al.* 2006, Dormann *et al.* 2009). See text for details.

In bipartite networks, *nestedness* is a central property that describes network structure. Studies evaluating beta-diversity have long recognized that species turnover among sites can be decomposed into nestedness and turnover components (Baselga 2010, Cardoso et al. 2014). When sites with lower diversity contain a subset of the species of sites with higher diversity, the beta-diversity is dominated by the nestedness component (Fig. 2A). Scaling up from pairwise habitat comparisons to the network level, a network is said to be nested when the communities of sites that have a few links (i.e. species) are a subset of the communities of sites with more links (Atmar and Patterson 1993). In a nested species-habitat network, the entire system will likely be affected if the most species-rich habitat or site is removed. In contrast, the removal of species-poor habitats that only interact with a few habitat generalists, is unlikely to have significant ripple effects.

In a bipartite network it is also possible to identify modules. A module comprises a set of habitat sites and species that interact more with each other than with other sites and species outside the module (Fig. 2B). *Modularity* measures the strength of division of a network into modules. Often, networks with a modular structure are expected to have a lower risk of collapse due to their buffering capacity to system perturbations (Dormann et al. 2017, Gilarranz et al. 2017). However, the loss of specific sites may also affect the associated species in the same module due to low redundancy. Hence, both *nestedness* and *modularity* can have profound conservation implications (Dormann and Strauss 2014).

A common way to assess those implications is to look at network *robustness*. The *robustness* of a network can be a key metric for conservation prioritization of high value sites and ecosystem management (Sole and Montoya 2001), as it is defined as the network resilience to the loss of nodes. For instance, simple simulations removing habitat randomly or in realistic sequences are one way to quantify community robustness to habitat loss (Fig. 2C). While particular species-habitat networks might be robust to random removals of habitats, they may also be highly sensitive to targeted habitat loss.

Understanding network *selectiveness* is central to assess the extent of habitat generalization (Blüthgen et al. 2006). An unselective network is characterized by having sites used proportionally to their size (green line in Fig. 2D), while selective networks are characterized by species using preferred sites, irrespective of site area (blue line in Fig. 2D). This metric can provide information about the consequences of different land-use change scenarios for species communities.

Species-habitat network analysis can also provide insights into the roles of specific habitat sites or species in the network. While some of these metrics can be derived from classic community ecology, the network approach enables scaling up to whole communities. First, the influence of one site upon another site can be assessed using *apparent influence* metrics (Muller et al. 1999). This index quantifies how much one habitat site contributes to sustaining the species present in another site (Fig. 2E). Interestingly, this index is not symmetrical (influence of node *a* upon *b* can be high, while the influence of node *b* upon *a* can be low) and more complex relations can be added, like adding time directionality in cases when the phenology of the habitat is known (e.g. flower phenology).

Another useful metric is node *strength* (Bascompte et al. 2006). This metric captures, for example, a single site’s importance taking into account how much the species depend on this site. A site can have high *strength* if it supports a high number of species with high dependency (i.e. specialist) on it (node *a* in Fig. 2F). Alternatively, sites that only host a few generalist species (node *b*) have low *strength* playing a minor role in the landscape (Collado et al. 2018).

We can also see the contribution of particular nodes to network level metrics like modularity or nestedness. The example of modularity is the most enlightening as modularity algorithms can also assess the role of each node in the network (Olesen et al. 2007). For example, using among-module *connectivity (c)* we can identify hub species connecting different modules (Fig. 2G). This can help to identify key-stone sites or species that can affect the robustness of the whole network. As for the whole network, habitat *generality* or *preference* can be also considered at the node level, using selectivity metrics (Fig. 2H) (Neu et al. 1974).

Overall, the characterization of nodes as individual habitat sites can be used to address questions regarding the extent to which particular network properties are related with ecological properties of the site (e.g. habitat quality, resources, area or isolation). Here, we have provided examples of a few commonly used metrics, while several comprehensive reviews of different metrics are available (Blüthgen et al. 2006, Dormann et al. 2009, Dormann and Strauss 2014). In Box 2, we present a worked example of species-habitat network using a published dataset.

#### Box 2. Analysing species-habitat networks: a worked example

Here, we illustrate a simple example of our approach by re-analysing a published dataset (Hill and Bartomeus 2016). The data comprises all bumblebee species sampled in multiple sites along 10 landscapes of 4 km^2^ (2 × 2 km) in Sweden. To exemplify how to apply common metrics, we will focus on a single landscape and build a species-habitat network. Even with a simple visualization as a bipartite network (Figure IIA), some ecological information can be obtained.

**Figure II.**
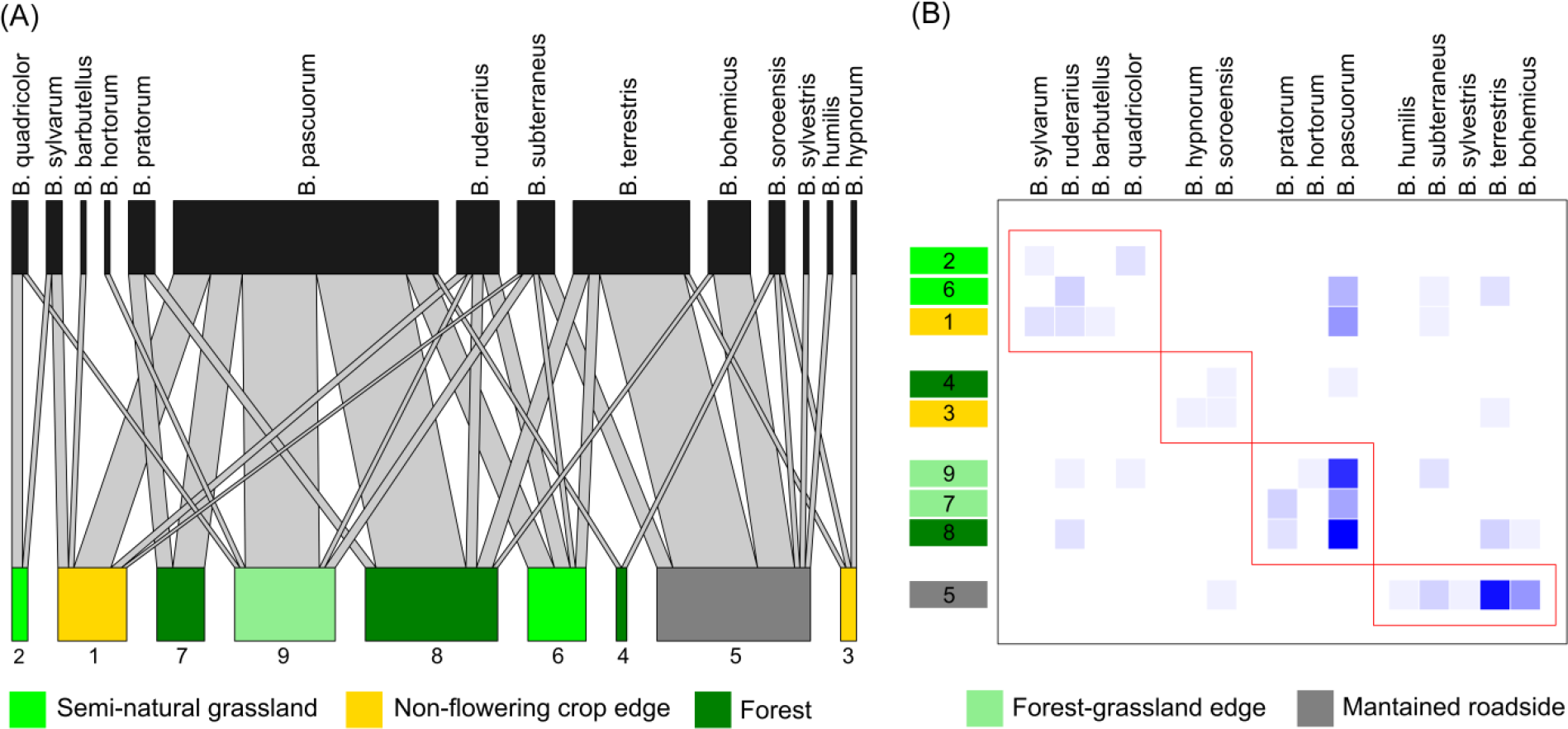
(A) Visual representation of the species-habitat network in one landscape where nine sites were sampled, and (B) plots showing the modules composing the network (red outline) with species abundance in each site (blue shading

For example, *B. pascuorum* is the most abundant species and is connected to most habitats, especially to semi-natural habitats and the maintained roadside is the most species-rich site. To facilitate conservation decision-making, we can calculate different metrics depending on the conservation aim. First, we show that this network is significantly more nested than expected by chance (observed NODF= 20.84, p< 0.001), i.e. species-poor sites tend to only host generalists that are also present in species-rich sites. If the aim is to protect the highest number of species with the minimum effort, a conservation strategy focusing only on the few most species-rich sites might be the best option. It is also possible to identify modules (Figure IIB), and calculate among-module (*c*) *connectivity* which in turn provides information about the role of each node in the network structure. Here, sites 6, 8 and 9 (*c* values close to 0.6) tend to act as connectors among different modules, and should thus be prioritized for conservation if the aim is to preserve the integrity of the network as a whole. To prioritize conservation, another option is to calculate site *strength* (*S*). The maintained roadside is the site with the highest *strength* (*S*= 4.36), because it hosts a large number of species and individuals that have a high dependence on this habitat. Finally, from a species perspective, we may be interested in habitat preferences. The *selectivity* (*d*’) index gives us information about the degree of habitat specialization of each species. With low values of *d*’, species such as *B. humilis* and *B. soroeensis* are among the most selective in this network. Since habitat specialists tend to be more vulnerable to extinction than generalists, these species should be the first included in conservation efforts (complete code to reproduce this and other analyses is available in the Supplementary Material).

## Implications for ecosystem management and policy

Land-use change is massively reshaping terrestrial ecosystems worldwide, and is recognized as a key driver of biodiversity loss with negative consequences on ecosystem functioning (Cardinale et al. 2012). An urgent question is to understand how to manage whole landscapes to maximize biodiversity conservation or ecosystem services delivery (Mendenhall et al. 2016). The flexibility of the proposed approach is that it can easily be applied across a range of ecological fields such as species conservation, habitat restoration, ecosystem services management, or invasion ecology (Memmott et al. 2007). Here, we provide four important research directions that could be addressed by adopting species-habitat networks:

a. *Conservation prioritization.* Conservation actions often face the trade-off between maximum protection of the environment and a limited budget. Site *strength* values in a landscape or in a protected area network can be used to prioritize which sites to conserve to maximize the biodiversity of any target taxon.
b. *Land-use change and community stability.* Conservationists often aim to achieve maximum biodiversity representation, without an explicit focus on the long-term stability. Seminal works (May 1972) and more recent studies (Thébault and Fontaine 2010, Gilarranz et al. 2017) on ecological networks have tried to use architectural patterns such as *modularity* to understand the mechanisms underlying the stability of communities. Similarly, we can investigate if certain species-habitat structures confer stability to the system in order to predict the *robustness* of species-habitat interactions to habitat perturbations.
c. *Maximizing biodiversity-based ecosystem services.* Landscape interventions to support ecosystem services often require the introduction of new habitats (e.g. hedgerows, mass-flowering crops) across a landscape. For instance, pollinators and pest control agents are known to be enhanced by the proximity to semi-natural areas (Ricketts et al. 2008, Schellhorn et al. 2015, Grass et al. 2016). Simulations using different crop configurations and green infrastructures can be used to maximize the positive *influence* among sites. For example, placing early mass flowering crops in the right configuration may maximize ecosystem service delivery, without imposing negative effects on natural habitats (Magrach et al. 2017).
d. *Impact of invasive species.* Landscapes are often invaded by alien species with a strong impact on native communities and ecosystem functioning. Here, the application of the species-habitat network will help to better understand the native community response to alien invasions across gradients of landscape composition and configuration. Incorporating a temporal perspective will elucidate how alien species move and use resources across the landscape. For instance, *modularity* or *selectivity* can provide key information on species spill-over and potential competition between natives and aliens.

The dichotomy of focal habitat versus the surrounding landscape overlooks the diversity of processes that characterise real-world landscapes. Species-habitat networks enable characterization of not only species or habitat-level dynamics, but also the emerging properties of those landscapes, going beyond the traditional landscape patch-mosaic model (Wiens 1995). By sampling multiple networks along relevant environmental gradients, these emerging properties can be used to test the effects of large scale drivers of global change upon ecosystem structure and stability (Schleuning et al. 2012). One key advantage of the application of the species-habitat network is that the scale of the derived ecological information will match the scale of landscape management interventions. The versatility, visualization power and easy interpretation of these networks will enable the application of the species-habitat network concept to a wide array of real-world problems concerning biodiversity conservation and ecosystem service enhancement at different spatial scales.

## Acknowledgements

We are grateful to P. Paolucci (Padova) for the drawings included in the figures.

## Data Accessibility

All code and data for creating the practical example included in the supplementary material is available at https://ibartomeus.github.io/hab-sp_ntw/demo.html.

